# EEG markers of vigilance, task-induced fatigue and motivation during sustained attention: Evidence for decoupled alpha- and beta-signatures

**DOI:** 10.1101/2024.10.16.618638

**Authors:** Simon Hanzal, Gemma Learmonth, Gregor Thut, Monika Harvey

## Abstract

Reduced vigilance can be captured in measures of attentional lapses in sustained attention tasks, but just how these lapses relate to task-induced fatigue and motivation to maintain optimal performance is unclear. We used the sustained attention to response task (SART) to induce fatigue, and manipulated motivation levels for the last block of the task in young and older participants (N = 34), while recording EEG to track electrophysiological markers of vigilance change, fatigue and motivation. Despite significant increases in subjective fatigue and mind wandering over 45 minutes, no vigilance decline was observed. However, the age groups differed markedly in their response strategies from the outset (adopting distinct speed-accuracy trade-off strategies) with faster/more erroneous responses in the younger and slower/more accurate responses in the older participants. The subjective rises in fatigue/mind wandering were coupled with an increase in pre-stimulus alpha-power, whereas the post-stimulus activity revealed two distinguishable beta signatures: a fronto-central topography as a marker of response strategy and a fronto-parietal distribution modulated by motivation *per se*. Our results thus show three distinct neural patterns underpinning the effects of fatigue, response strategy and motivation and suggest a (motivational) cognitive control mechanism behind resetting of performance decrement, independent of persistent fatigue.

## Introduction

Sustaining attention requires a constant, self-directed maintenance of vigilance (Robertson and Garavan 2004) both across various daily activities (Roach et al. 2012; Massar et al. 2018; Walker and Trick 2018), and bespoke experimental tasks (Head and Helton 2012; Reteig et al. 2019). Continuously engaging in sustained attention tasks can result in changes in performance (Stoll et al. 2016; Reteig et al. 2019) that have been characterised as the *vigilance decrement* (Robertson and Garavan 2004; Oken et al. 2006). However, the onset of a vigilance decrement will vary across experiments and studied populations, with some studies pointing to a continued ability of participants to concentrate and maintain focus during sustained attention tasks (Nakagawa et al. 2013; Lara et al. 2014), while others suggest impairments in accuracy and reaction times with increasing time-on-task (Smit et al. 2004; van Schie et al. 2012; Pershin et al. 2023).

Differences in motivation, both between-and within-participants, may account for some of this variability. For instance, it has been suggested that motivation can be a key influence in the reappraisal of task strategies (Earle et al. 2015; Gilsoul et al. 2022), which can lead, through the self-regulatory mechanism of attentional effort (Sarter et al. 2006; Stoll et al. 2016), to improved behavioural measures in sustained attention tasks (Oken et al. 2006). This was tested by Reteig et al. (2019), who rekindled participants’ motivation after 60 minutes spent on a sustained attention task, by offering an additional monetary reward if they managed to out-perform 65% of the other participants in the final part of the experiment. Although Reteig et al. (2019) found that the motivational manipulation restored the vigilance decline to some extent, this was not reflected in the tested EEG measures of attentional control, except for variability in a neural theta-response. Hence, a neural link of the effect of motivation on vigilance change remains tentative (see also (Awh et al. 2012)). In the present study, we sought to re-examine the role of motivation in vigilance decrements by manipulating motivation in the course of a sustained attention task while recording its EEG markers (similarly to (Reteig et al. 2019)), and further extending the investigation into the neurophysiological basis of fatigue.

Performance declines in sustained attention tasks are often induced by mental fatigue, which can be characterised as a dynamic, time-dependent, state-like response (Ishii et al. 2014; Völker et al. 2016; Wylie and Flashman 2017; Reteig et al. 2019), typically resulting in difficulty concentrating and/or enhanced mind-wandering (Martínez-Pérez et al. 2023). The exact mechanisms driving mental fatigue are still unclear (Kuppuswamy 2022), but are likely caused by either a depletion of cognitive resources, or by monotonous tasks leading to a disengagement of the sustained attention networks (Gergelyfi et al. 2015). Notably, previous work highlights that the effect of fatigue on vigilance decrement may be decoupled from the effect of motivation (Gergelyfi et al. 2015) and hence taking into account fatigue in the study of neural markers of vigilance change, in addition to motivational factors, is relevant. Unlike (Reteig et al. 2019), who used a sustained attention-style task where the goal was to detect rare targets in an oddball paradigm (and where typical errors were omissions), we used the sustained attention to response task (SART) (Robertson et al. 1997; Weinstein 2018) in which failures in sustained attention manifest in commission errors to infrequent no-go stimuli. Performance changes in the SART are influenced by response strategy and chronobiology (Lara et al. 2014; Wilson et al. 2016; Dang et al. 2018), can capture lapses into a less attentive state (Wilson et al. 2016), as well as task monotony (Head and Helton 2012), so the mapped effects extend well beyond the task itself (Smit et al. 2004). Prolonged versions of the SART therefore represent an optimal method for exploring the relationship between vigilance decrements and potential differences in strategy linked to motivation and fatigue.

In terms of neural substrates and/or correlates of the vigilance decline, both the relevant anatomical networks and some of its potential neural markers have been identified. The vigilance network is responsible for sustained attention (Milyavskaya et al. 2021), as well as executive function (Holtzer et al. 2011), exerting attentional control over the incoming visual stimuli (Posner and Dehaene 1994) via modulation of early visual centres and network interactions needed to carry out the task (Corbetta et al. 2002; Clayton et al. 2015). It has also been linked to task-induced fatigue (Shen et al. 2016) and brain activity potentially associated with this network has been observed to change over the course of a task (Macdonald et al. 2011; Benwell et al. 2019; Li et al. 2020). Accordingly, activity changes in the vigilance network should track its gradual disengagement (Johansson and Rönnbäck 2013; Ishii et al. 2014) and thus make the network a viable focus for the study of neural changes due to the vigilance decrement.

In EEG research, activity of the vigilance network has been broadly characterised as occurring in the oscillatory alpha frequency band (Clayton et al. 2015; Sadaghiani and Kleinschmidt 2016)) extending over frontoparietal brain areas (Corbetta and Shulman 2011; Clayton et al. 2015), although oscillatory activity pertaining to control extends beyond the alpha-band. Changes in beta-band activity have been associated with the regulation of attentional effort during time on task, in addition to motor preparation (Liu et al. 2010; Stoll et al. 2016; Li et al. 2022). Thus, analyses of oscillatory signals focusing on the alpha and beta band (Liu et al. 2010; Stoll et al. 2016; Benwell et al. 2018; Arnau et al. 2021; Li et al. 2022) should contribute to a better understanding of the neural processes underlying vigilance changes (Martínez-Cañada et al. 2023) and clarify their contribution to fatigue and motivation.

Aside from oscillations, changes in the P300 ERP component have been associated with subjective fatigue (Krigolson et al. 2021), as well as time on task in sustained attention paradigms (Hart et al. 2012; Guo et al. 2016), and ageing (Kaufman et al. 2016). We thus mapped the P300 ERP-changes across the factors of interest (time on task, fatigue, the motivational manipulation), as well as age group.

Finally, it is well documented that there is large inter-individual variability in SART performance across the population (Vallesi et al. 2021; Hanzal et al. 2024). In particular, different age groups tend to adopt either an accuracy (Dang et al. 2018; Reteig et al. 2019) or speed-based (Lara et al. 2014; Statsenko et al. 2020) strategy. These strategies are then prone to change (for example, switching from an emphasis on achieving high accuracy, to responding faster) during the task (van Schie et al. 2012). This could reflect differences in the underlying levels of fatigue, as suggested by surveys of general population fatigue (Gilsoul et al. 2022; Yoon et al. 2023), or instead an age difference in motivational levels (Ryan and Campbell 2021; Carr et al. 2022).

In brief, we investigated in this preregistered study the behavioural and EEG measures of the vigilance decrement and increased fatigue in young and older participants as a function of time-on-task during prolonged (45 minutes) SART performance. We induced high levels of motivation in half of the participants during the final experimental block to investigate whether motivation could improve performance and change EEG markers of fatigue and tested for differences between age-groups. Our results reveal three, dissociated, frequency-specific EEG signatures of increased fatigue/mind wandering with time-on-task, of response strategies per age-group and of motivation, on the backdrop of maintained task performance (no vigilance decline observed). We speculate that these signatures may contribute to offset performance declines over time-on-task.

## Materials and Methods

### Participants

The hypotheses, design and analysis plan were pre-registered prior to data collection and can be accessed via the Open Science Framework (https://osf.io/y2vgc/). The study was approved by the University of Glasgow College of Science and Engineering Ethics committee. A total of 41 healthy adults aged between 18 and 87 years old were recruited from the University of Glasgow subject pool and the local area and were given monetary compensation for their time. The study was approved by the University of Glasgow College of Science and Engineering Ethics committee (Approval number: 300210156). Written consent was obtained from all participants. Participants were balanced for gender and were asked to report any existing medical conditions, eye-sight correction and medications which might impact their performance. Seven participants were excluded due to excessive noise and artifact in the EEG signal. The final sample consisted of 34 participants (F = 16) split into two groups based on age: young (n = 18, F = 9, mean age = 22.61, SD = 1.85, range = 20 - 26) and older adults (n = 16, F = 7, mean age = 66.50, SD = 8.45 years, range = 55 - 87). Two participants were left-handed, one was a smoker and all participants reported low to moderate caffeine consumption (estimated mean units per day = 1.31, SD = 1.12, range = 0-4), corresponding to the maximum recommended daily dose of 400mg of caffeine (Mitchell et al. 2014). They also reported an average of 7.34 hours of sleep per day (SD = 0.85, range = 6-9). All young participants were enrolled university students and the older group had similar levels of tertiary education (n = 6, 37.50%) compared to the UK average for their age group (39.60%) (OECD 2023).

All participants were screened for cognitive difficulties using the Montreal Cognitive Assessment test (MoCA; (Nasreddine et al. 2005)), reflecting scores representative of a healthy population (Borland et al. 2017) in both young (mean score = 28.28, SD = 1.49, range = 26 - 30) and older adults (mean = 25.81, SD = 2.74, range = 22 - 30). A short (3 minute) computerised visual screening assessment was administered at the beginning of the session to exclude potential visual pathologies. The task was adapted from a similar experiment investigating lateralised visual attention in both young and older groups (Learmonth et al. 2017) and shortened to 32 trials. A Welch’s t-test identified no between-group differences in target detection within the visual regions where the SART stimuli were to be presented, t(32) = 1.45, p = 0.163.

### Subjective Measures

Changes in state fatigue were assessed by the Visual Analogue Scale for Fatigue (VAS-F). The VAS-F measures 18 items across two subscales (fatigue = 13 items and energy = 5 items), with scores of 0 = low fatigue to 100 = high levels of fatigue. It has excellent test-retest reliability of α = .93 and α = .91 for the two scales, respectively (Lee et al. 1991). As in Hanzal et al., (2024), two items on the scale were replaced with synonyms: “worn out” was changed to “drained”, and “bushed” to “run down” to avoid repetitiveness and dated language. The spontaneous subscale from the Mind Wandering measure (Carriere et al. 2013), comprising 4 items on a 7-point Likert scale was administered to measure changes in mind-wandering during the experiment.

### Sustained Attention to Response task (SART)

The study used a custom version of the SART (Robertson et al. 1997) (Figure 1B). In each trial, participants were instructed to maintain fixation on a centrally presented cross and attend to a numeric stimulus (0-9) presented at an angular distance of 1° for 250ms. The fixation reappeared for a variable duration of 3000-4000ms before progressing to the next number. The stimuli were black on a white background and presented using a 21-inch CRT monitor (Samsung, SyncMaster 1100MB) with a screen resolution of 1024 x 768 pixels and a refresh rate of 100 Hz. Participants were seated 60 cm from the screen, maintaining horizontal eye level with the centre of the display. Participants were instructed to click the left mouse button in response to all numbers that appeared (go trials), apart from 3 and 6 (no-go trials). The participants did not receive any feedback about their individual response times or accuracy. The stimuli were pseudo-randomised to ensure equal frequency and random distribution throughout the experiment.

**Figure 1.**
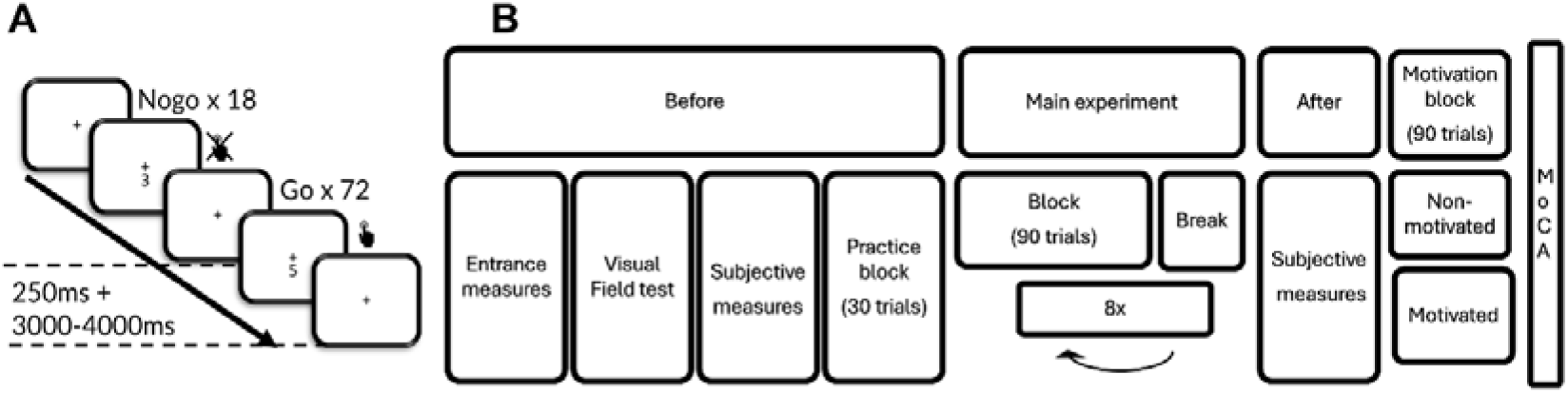
**(A)** Outline of the SART task. **(B)** Outline of the experimental procedure. Before the task, participants completed the entrance demographic information, the visual field test and were administered the subjective measure questionnaires assessing state fatigue and mind wandering (VAS-F, MW-S). For the main experiment, participants performed 8 blocks of the SART task, allowing tracking of vigilance over time. After completing the experiment, the participants again completed the subjective measure questionnaires (VAS-F, MW-S) and proceeded to the final motivational block of SART performance. The MoCA was administered at the end to assess potential presence of cognitive impairments in the sample.

### Procedure

The experimental procedure is outlined in Figure 1. Participants were provided with an information sheet and gave written, informed consent to take part. They then provided basic demographic information and reported visual deficiencies, as well as any other relevant medical conditions. A 64 Ag/AgCl BrainCap (BrainProducts, Gilching, Germany) were fitted according to the international 10/20 system (American Electroencephalograpic Society, 1991), including two horizontal electro-oculographs, and impedance reduced to <25KΩ using Signa gel. Participants then completed the VAS-F and MW-S and a brief visual field test to map visual acuity. They then underwent a SART practice session of 30 trials and were given general feedback about their accuracy on the practice block. In the main experiment, participants completed eight blocks of the SART (90 trials, 72 go, 18 no-go trials, random presentation), each lasting five minutes and 20 seconds, with self-paced breaks between each block, followed by the VAS-F and MW-S to record post-task subjective states. Continuous EEG data was recorded at a 1000Hz sampling rate. The first eight blocks of the SART were followed by one further, unannounced, block of 90 trials (Block 9) to manipulate motivational state. Half of the participants were randomised into a motivated group and were instructed to try to maximise their performance on the last block (no guidance was given regarding potential strategies to achieve maximal performance). They were informed that the participant with the best performance during this block would receive an additional bonus of £50. The other half were only informed that the experiment includes an additional block and asked to undertake the block with the same instructions as the previous blocks. Finally, the participants were screened for mild cognitive decline using the MoCA and debriefed.

### Analyses

#### Behavioural analyses

All behavioural analyses were carried out in R (R core team 2020) using the packages ‘tidyverse’ (Wickham et al. 2019), ‘psych’ (Revelle 2023), ‘moments’ (Komsta and Novomestky 2022), ‘readxl’ (Wickham et al. 2019), ‘broom’ (Robinson et al. 2024), ‘ez’ (Lawrence 2016), ‘lmerTest’ (Kuznetsova et al. 2020), ‘lme4’ (Bates et al. 2015), ‘emmeans’ (Lenth et al. 2024). Further packages used for graphical depiction were: ‘ggpubr’ (Kassambara 2023), ‘viridis’ (Garnier et al. 2023) and ‘Cairo’ (Urbanek and Horner 2023). Performance analyses of the effect of age-group and time on the chosen metrics relied on an examination using randomised block- and participant-level modelling. The lmer function of the ’lme4’ package (Bates et al. 2015) was used to construct corresponding random mixed effects model and to fit individual participant and block slopes and intercepts, with the lmerTest package (Kuznetsova et al. 2020) to estimate p-values.

#### Electroencephalography (EEG) analyses

Analysis of the EEG data was undertaken using the EEGLAB (Delorme and Makeig 2004) and FieldTrip (Oostenveld et al. 2011) toolboxes for Matlab. The continuous data was first detrended to remove drifts introduced by instrumental and physiological noise, alongside various baseline shifts. A Hamming-windowed FIR filter was then applied within the 2-45Hz frequency range, followed by re-referencing to the average signal. Independent component analysis was then run using ICLabel (Pion-Tonachini et al. 2019). The following threshold criteria were applied to identify components for automatic rejections: 1) Components that had <0.05 likelihood of brain origin and 2) Components that had >0.8 likelihood to be one of either an ocular artefact, muscle artefact, heart artefact, line noise or channel noise. All components labelled as “Other” were visually inspected and rejected if they appeared similar to standard artefact components. This semi-automated correction method led to the rejection of a mean of 25.97 components (of a total of 64 available components per person, SD = 4.25, range = 20 - 37). A further 1.18 (SD = 1.40, range = 0 -3) components were manually rejected in each dataset and 0.24 (SD = 0.55, range = 0 - 2) preserved from automatic rejection due to incorrect ICLabel classification. The same analytical steps were then performed on the raw datasets with a lower filtering threshold of 0.5 - 40Hz, the original ICA weights were re-applied and components removed. Upon inspection of the signal, known noisy electrodes were interpolated (mean per participant = 0.76, SD = 0.96, range 0 - 4).

Data for analyses were selected based on the following steps:

1. Participants with insufficient neural data as outlined in data pre-processing were not included in behavioural analysis.
2. Trials with trigger information missing due to failure of transition of an event signal were identified and removed.
3. Commission trials were removed based on deviant reaction time. Firstly, block and group mean and standard deviation in reaction time was computed. Blocks were considered separately due to the expected time effect and groups were used because of the effect of cognitive strategy. Then trials rising above two standard deviations of the mean were removed as attention lapses and trials two standard deviations below the mean as anticipation error (Kiesel et al. 2008).
4. Participants were also removed if they exhibited any of two identified erroneous strategies in any of the 8 main experimental blocks: a) responding to all trials at chance level (> 80% go stimuli correct and < 20% no-go stimuli correct), or b) withdrawing the response for all trials at chance level (< 20 % go stimuli correct and >80% no-go stimuli correct) in any of the eight experimental blocks.

Trials thus rejected (based on behaviour) were also identified and removed from the EEG analysis. Further trials were identified for removal based on eye-inspection of the signal, detecting ICA-nonremoved artefact, leading to a rejection of 4.60% of trials (SD = 2.70, range = 0.00 – 18.19%). After initial cleaning, the data was re-epoched for pre-stimulus and task-related analysis. Time-frequency analysis was performed using a transformation based on multiplication in the frequency domain method as specified in the ft_freqanalysis Fieldtrip function (Oostenveld et al. 2011), and a Hanning taper was applied to the data. The frequency range of interest was defined as 2 to 40 Hz with a 1/3 Hz frequency step. The number of fixed cycles per wavelength was set to 6. For all permutation testing, spatial neighbours for each electrode were defined as those being approximately 5cm distant (Maris and Oostenveld 2007). The maximum possible number of permutations (up to 3000) was undertaken for each test. To investigate whether neural patterns differed across conditions of interest, a permutation test was run on all channels over the whole epoch (-1500ms to 1500ms) applying relative baseline correction and transforming the data to decibels.

A separate data cleaning procedure led to the selection of different participant data for the motivational group, leading to 33 (group 2 = 13). This is explained by a case-by-case basis of inclusion of participants based on the overall neural data quality which fluctuated between the two parts of the experiment, with a final overlap of 30 participants between both parts. The motivational block data was notably noisier, due to a likely higher proportion of agitation and motion artefacts introduced by the motivational manipulation.

The procedure for analysing the pre-stimulus signal was guided by the recent decomposition of the signal into periodic and aperiodic components (Donoghue et al. 2020) which have distinct associations with tasks and participant populations (Donoghue et al. 2022), e.g. ageing (Turner et al. 2023) and pathology, as in developmental dyslexia (Turri et al. 2023). This was undertaken using the ‘mne’ python package (Gramfort et al. 2013). The time-frequency signal was examined through spectral parameterization in trial-averaged time-frequency spectra for each channel and each participant in blocks 1 and 8, using an implementation of the aperiodic analysis through the ‘specparam’ python package (Donoghue et al. 2020). The resulting aperiodic 1/f fit was removed from the power spectrum using subtraction. The corrected ‘periodic’ spectrum was permutation-tested using the permutation_cluster_1samp_test and permutation_cluster_test functions as implemented in the ‘mne’ python package (Gramfort et al. 2013), by supplying first and last block difference matrices per each participant. The t-statistic significance threshold was manually determined using the ppf function from the ‘scipy’ package. The adjacency sparse matrix was obtained using the native easycap 64 channel layout derived from the MATLAB ‘fieldtrip’, and extended to include N-1 and N+1 neighbours along the frequency dimension using the combine_adjacency function implemented in the ‘mne’ package.

## Results

### SART Task Performance

Across the two groups, the young adults showed faster responses in terms of reaction times (mean = 429.45ms, SD = 50.37ms, range = 328.88ms – 577.67ms) (Figure 2A1) but lower accuracy due to elevated commission errors (i.e. a failure to withhold responses; mean = 24.46%, SD = 13.19%, range = 0%-61.11%) (Figure 2B1), while the older group showed a reversed pattern, with slower reaction times (mean = 567.66ms, SD = 95.18ms, range = 429.99ms – 923.98ms) but fewer commission errors (mean error = 7.72%, SD = 7.48%, range = 0-33.33%) (see the same Figure 2A1 and Figure 2B1). Omission errors (missed targets) showed floor effects both in young (0.21%, SD = 1.15%, range = 0 – 12.5%) and older adults (mean error = 0.80%, SD = 2.03%, 0 – 13.89%) alongside a heavy skew, so omission errors were not analysed further.

**Figure 2.**
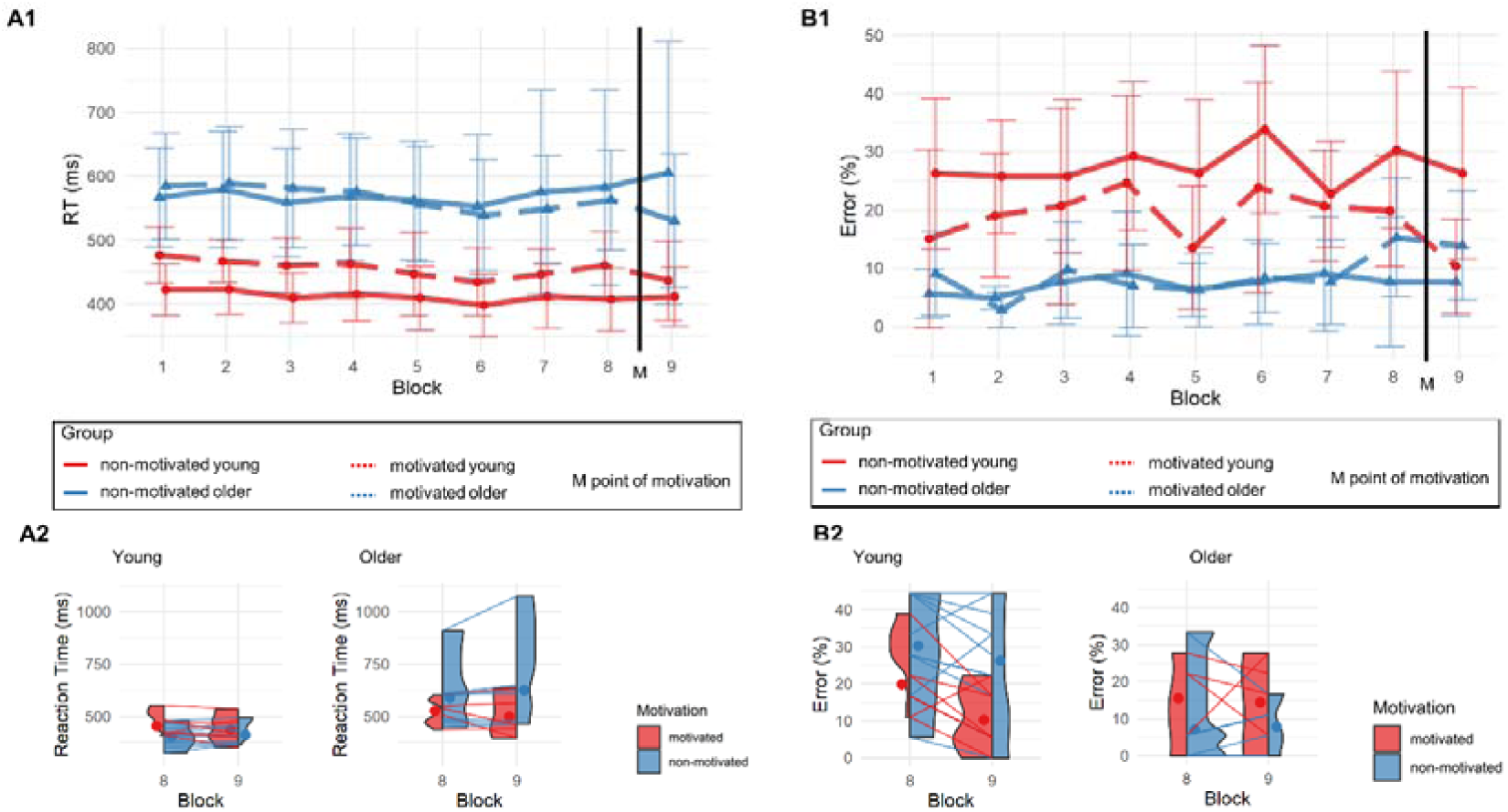
Behavioural Performance. (A1) Mean reaction time and (B1) commission error in both experimental groups (older in red, young in blue) across time-on-task (experimental blocks 1-8) and a final experimental block (block 9) with/without motivational manipulation (motivated, nonmotivated subgroup). (A2) Difference in reaction times and (B2) commission error induced by the motivational manipulation in both age groups (young, older) as inferred from comparisons between blocks (8, 9) in both the non-motivated group (blue) and motivated group (red).

To track possible vigilance decrements over time, we tested the effect of time-on-task (experimental blocks 1-8) and age group (young vs old) on commission error using a random mixed effects model. The effect of time-on-task was not significant [t(32) = 0.43, p = .159] but lower error rates were confirmed in older participants as opposed to the young group [t(32) = 15.77, p < .001] with no interaction [t(32) = 0.36, p = .718]. An identical model was run to analyse reaction times. The effect of time-on-task was significant, indicating a slight reduction in reaction times across time [t(32)= 0.01, p = .049] likely due to learning effects, alongside a main effect of age group, showing faster reaction times in young participants as opposed to the older group [t(32) = 0.23, p < .001], with no interaction [t(32) < 0.01, p = .775].

We then tested the effects of the motivational manipulation using a three-way ANOVA with the factors block (block 8 vs 9), age group (young, old) and motivational group (non-motivated, motivated) separately for reaction times and commission errors. The motivational manipulation consisted of instructing half of the participants after completion of block 8 that they can earn more money if they outperform their fellow participants in the last block (block 9), while the other half were simply told that there is one final block to be completed (motivated vs unmotivated subgroup).

The analysis of the reaction times showed that there was no effect of block [F(1, 26) = 0.37, p = .550, η² < .001] and no effect of motivation [F(1, 26) = 0.18, p = .676, η² < .001], but the interaction between block and motivation was significant with a small to medium effect size [F(2, 26) = 7.74, p = .010, η² = .016]. Confirming the analysis above, the older group had slower reaction times than the young group [F(1, 26) = 16.17, p < .001, η² = .418]. Post-hoc analyses were conducted using paired sample t-tests to further explore the interaction of block and motivation. These revealed that in the non-motivated group, reaction time increased between blocks 8 and 9 [t(17) = -2.47, p = .024], while remaining stable in the motivated group [t(11) = 1.65, p = .128], hence suggesting that the motivational manipulation had an effect on reaction time. See Figure 2A2. No other interaction was significant.

The analyses of the commission errors showed that there was no effect of block [F(1, 26) = 1.535, p = .226, η² = .045] and no effect of motivation [F(1, 26) = 1.53, p = .226, η² = .04]. The results also showed that the older adults were more accurate than the younger group [F(1, 26) = 10.30, p = .004, η² = .236]. Unlike for the reaction times results, there was no interaction between block and motivation, hence suggesting that there was no effect of the motivational manipulation on commission errors. There was a significant interaction between age group and motivation [F(2, 26) = 4.77, p = .015, η² = .172] but this interaction was not further explored due to the absence of an interaction with block. This 2-way interaction is picking up on a difference between motivated vs non-motivated young participants independent of block (despite random allocation) that is not seen in the older participants and is almost certainly reflecting a chance effect despite the random allocation of participants to groups. See Figure 2B1 for an illustration of the interaction that is seen throughout all blocks. No other interaction was significant.

### Subjective levels of fatigue and mind wandering

Subjective fatigue levels were assessed before and after task performance (pre block 1 and post block 8). From the beginning, the older group (mean = 427.56, SD = 201.22, range = 104 - 755) showed lower baseline fatigue scores than the young group (mean = 588.89, SD = 257.61, range = 118 – 1054) [t(32) = 2.05, p = .049, see Figure 3A]. Likewise, from the beginning, older adults showed lower mind wandering scores (mean = 3.70, SD = 1.61, range = 1.25 – 6.50) than the younger group (mean = 4.07, SD = 1.71, range = 1 - 6) [t(32) = 2.69, p = .010, see Figure 3B].

**Figure 3.**
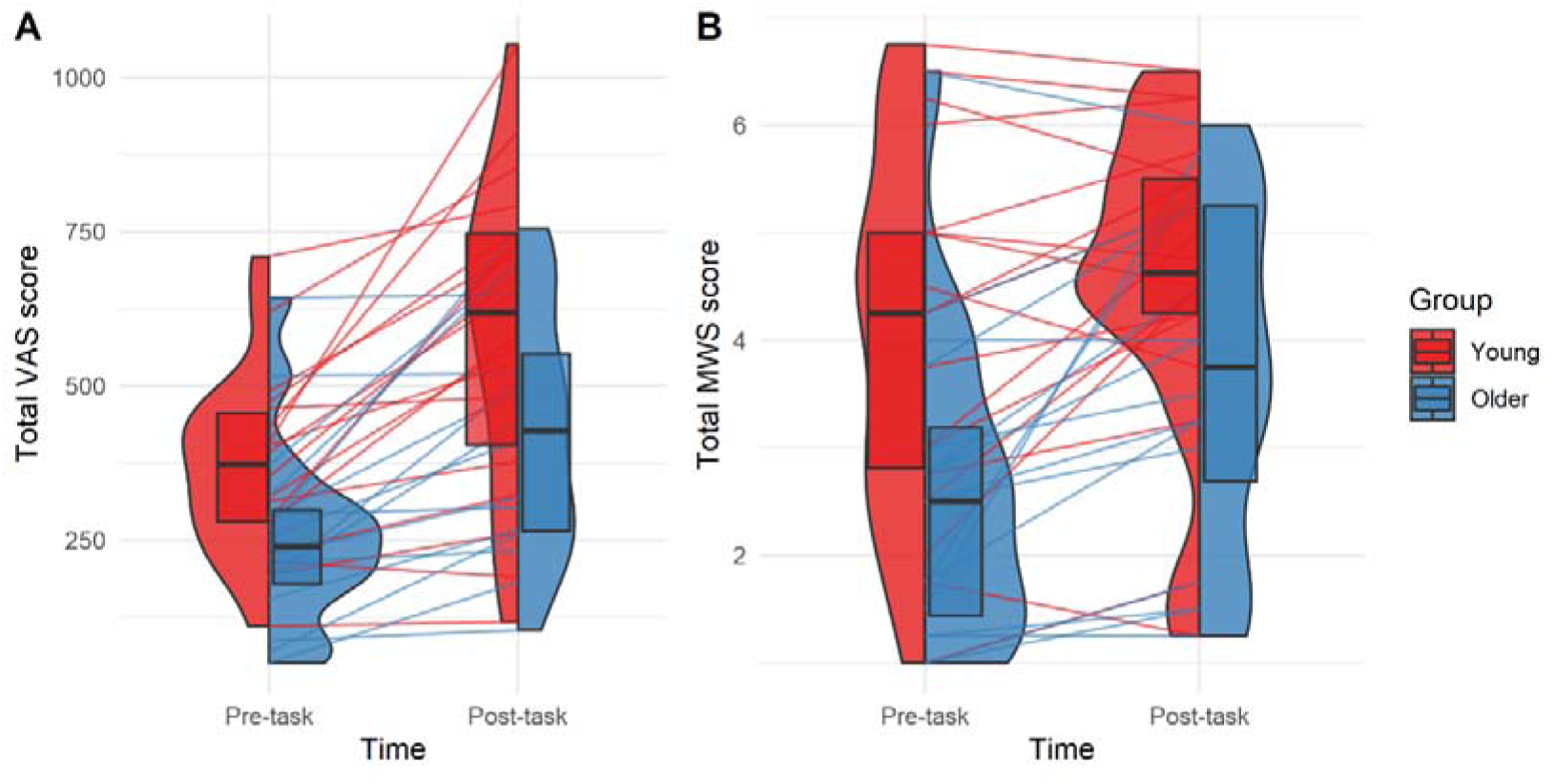
Subjective measures. (A) Scores of the visual analogue scale for state fatigue and (B) mind wandering before and after the experiment (Pre-task, Post-task) for each group (young in red and older in blue), with individual participant total scores, as well as overlaid boxplots with overall medians and quartile ranges.

Comparing pre-task with post-task subjective scores (see Figures 3A and 3B), we found that both the young (mean change = 217.06, SD = 182.29, range = - 25 - 646) and older group (mean change = 169.31, SD = 149.58, range = 3 - 485) showed a rise in their subjective fatigue levels with time-on-task. Likewise, both the young (mean change = 0.53, SD = 1.00, range = -0.75 – 2.50) and the older group (mean change = 1.11, SD = 1.28, range = -0.50 – 4.25) had an increase in mind-wandering scores.

In a 2x2 mixed ANOVA testing the effect of time (before, after), age group (young, older) and their interaction on the subjective fatigue scores, we found a large effect of time [F(1, 32) = 45.75, p < .001], showing that subjective fatigue scores had increased by the end of the main experiment. There was also a small difference between the groups, indicating that the young adults had slightly higher fatigue scores than the older group [F(1, 32) = 5.03, p = .032]. However, there was no significant interaction between group and timepoint [F(2, 32) = 0.69, p = .414].

Similarly, in a 2x2 mixed ANOVA testing the effect of time (before, after), age group (young, older) and their interaction on mind wandering, there was a large effect of time, explained by higher mind wandering scores by the end of the experiment [F(1, 32) = 16.80, p < .001]. There was again a main effect of age group, with the young group having higher mind wandering scores than the older group [F(1, 32) = 5.61, p = .025]. However, there was no interaction between group and timepoint [F(2, 32) = 2.20, p = .147]. To test for a link between mind wandering change and subjective state fatigue change, a linear regression was fitted to the paired observations for each participant, but the model was not significant [F(1, 32) = 2.42, R^2^ = 0.041, beta < 0.01, p = .130].

### Electroencephalography (EEG) results

The EEG-analyses aimed at identifying the brain oscillation markers of the above performance differences and the reported subjective changes, across our experimental factors. Due to the absence of the vigilance decline, we first sought to identify the EEG-changes with time-on-task as a possible marker of increased fatigue/mind wandering, or alternatively, of enhanced effort to maintain stable task performance despite the reported subjective increase in fatigue. In a second step, we examined co-variations with age-group and the motivational manipulation. Finally, we tested for correlations with behavioural measures. This was to disentangle EEG-markers (i) of fatigue/mind wandering *per se*, which should show changes both across age-group (enhanced fatigue in the younger as compared to the older group) and time-on-task (increasing fatigue across blocks in each group). II) to disentangle EEG markers of the difference in response strategy in the older vs the younger participants (deployment of effort more towards motor control/accuracy versus response speed), and iii) of the effects of the motivation manipulation. Our results reveal EEG-markers (i) of fatigue/mind wandering in pre-stimulus alpha-oscillations over centro-parietal locations, (ii) of qualitative differences in response strategy/ effort deployment in post-stimulus, task-related beta-desynchronization/rebound over fronto-central sites, and (iii) of motivational manipulation in a distinct, fronto-parietal beta-signature, as outlined below.

### Pre-stimulus EEG oscillations: alpha-signals increase with time-on-task, differ by age-group and are amenable to motivational manipulation

We first investigated potential oscillatory markers of time-on-task. We argue that these likely reflect either correlates of enhanced fatigue/mind wandering or enhanced effort to execute the task at stable performance levels, given the absence of a performance/vigilance decrement over time-on-task, despite increased fatigue (from block 1 to 8). To identify the effects of time-on-task on oscillatory activity, we first compared activity in the pre-stimulus window (-1100 to 0ms) across a broad spectrum of frequencies (3-40Hz, including the alpha- and beta-band) between experimental blocks 1 and 8, whereby we decomposed the full pre-stimulus spectrum per experimental blocks and group (Figure 4A) into its separate periodic (Figure 4B) and aperiodic components (Figure 4C) (Donoghue et al. 2020), and ran comparisons using cluster-based permutation statistics (Maris and Oostenveld 2007). We then examined whether the markers of time-on-task (Figure 4D) co-vary with other possible contributors to task performance/vigilance, namely age-group and motivation (Figures 4E-F).

**Figure 4.**
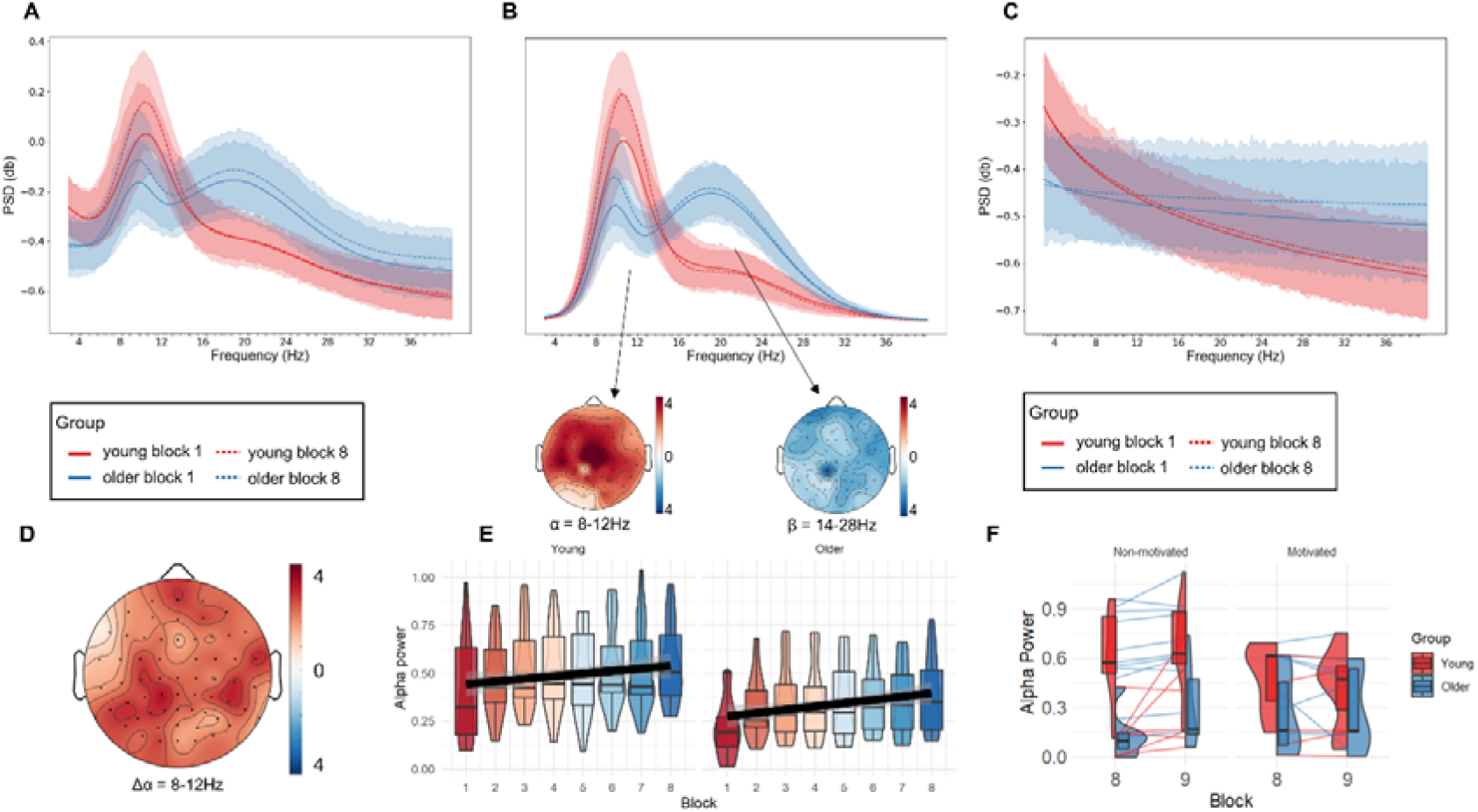
Time-on-task related pre-stimulus activity. Spectrograms for (A) total measured power, (B) the periodic signal after adjustment from the aperiodic component and (C) the aperiodic component, represented across Time-on-task (block 1 vs 8: solid line vs dashed line) and the young (in blue) and older group (in red). Transparent areas represent 95% confidence intervals. Note in (B) that the largest periodic spectral differences are observed between age groups in the alpha-band (left topography: alpha older < younger) and the beta-band (right topography: beta older > younger group). (D-E) Time-on-task of periodic alpha (8-14Hz) activity: The panels illustrate the alpha-increase with time-on-task in terms of (D) its topography (from block 1 to block 8) collapsed across age groups, and (E) its evolution across experimental blocks 1-8 for each of the age-groups separately (young vs old: left vs right). The plot illustrates the variability of the signal in individual participants, as well as the median values and quartile ranges with an overall positive linear trend fitted separately for each group. (F) Motivational manipulation and alpha-band activity: Plots denote periodic alpha-band changes from blocks 8 to 9 (with motivational manipulation), per each of the groups (younger: red vs. older: blue/ motivated: solid vs. non-motivated: dashed lines). The motivational manipulation influenced alpha-band activity by preventing further alpha-increases from block 8 to 9 in the motivated but not the non-motivated group, independently of age-group. Participant-level changes are depicted alongside median values and quartile ranges.

Examining the effect of time-on-task through cluster-based statistics of the periodic, pre-stimulus activity in experimental blocks 1 versus 8, revealed one positive cluster (cluster statistic = 3391, p = .007). This indicates a rise in power across time-on-task in the alpha frequency band (8-14Hz) over the majority of channels with a maximum in centro-parietal locations (see Figure 4D).

To characterise the alpha-increase with time-on-task over all experimental blocks 1 to 8 and to further test for possible co-variations with age-group differences, and motivational effects, we extracted the periodic alpha signal (8-14Hz band) of the pre-stimulus epoch from the electrode with the highest cluster t-statistic (CP1) across all blocks 1-9.

Figure 4E illustrates the periodic alpha-changes over time-on-task (blocks 1-8) per age-group. Analysis of the factors time-on-task (block 1-8) and age-group (young, old) using a random mixed effects model (with randomised participant- and block-level effects) revealed a significant increase in the alpha-signal across blocks [t(32) = 3.06, β = 0.10, p = .004] with the signal being elevated in the young group as compared to the older group [t(32) = - 2.29, β = -0.14, p = .030], but with no interaction [t(32) = -.217, β > -0.01, p = .830]. Note that directly examining the effects of age-group on the pre-stimulus, periodic component using cluster-based statistics (older vs younger participants) confirmed the age-effect in the alpha-band (alpha old < young), revealing one large negative, alpha-band cluster (highest cluster statistic = 9546, P = .001; see Figure 4B for cluster map), alongside a large positive cluster (cluster statistic = 8464, p = 0.001) in the adjacent beta band (14-30Hz). The latter showed elevated beta-activity in the old relative to the young group (see Figure 4B, spectrograms and cluster map).

Figure 4F illustrates the pre-stimulus alpha-signal that is sensitive to time-on-task (extracted from CP1) with regard to the co-variation with motivational manipulation, namely across block 8 and 9, separately for the motivated and non-motivated groups. Conducting a 2x2x2 ANOVA with the factors motivation group (motivated, non-motivated), age group (young, older) and block (block 8, motivational block 9) revealed no main effects of motivation on this alpha-signal ([F(1, 26) = 1.00, p = .326, η² = .034]), but an interaction between motivation x block (F(1, 26) = 5.00, p = .034, η² = .018). In addition, the young group had a consistently higher alpha-signal than the older group confirming the analysis above [F(1, 26) = 12.94, p = .001, η² = .310], with no difference between blocks [F(1, 26) = 3.42, p = .076, η² = .013]. No other interaction was significant. Post-hoc analyses were conducted using paired t-tests to further explore the interaction of motivation group and block. These showed that the alpha-signal of the non-motivated group continued to increase between block 8 to 9 [t(17) = 2.48, p = 0.024], while the motivated group showed no further change in this signal [t(11) = .616, p = .551].

Analysis of the aperiodic component (exponent and intercept) using 2x2 ANOVAs with the factors time-on-task (block 1 vs 8) and age-groups (young, older) revealed significant group-effects for both the exponent (steeper exponents in the young group [F(1, 32) = 13.99, p < .001, η² = .265]) and the intercept [higher intercepts in the young group [F(1, 32) = 8.57, p = .006, η² = .200]) but no effects of time-on-task [exponent: [F(1, 32) = 0.46, p = .503, η² = .003], intercept: [F(1, 32) = 0.28, p = .597, η² < .001]] nor any interaction (exponent: [F(2, 32) = 0.68, p = .416, η² = .004], intercept: [F(2, 32) = 0.50, p = .483, η² = .001]).

In summary, these results reveal that periodic, pre-stimulus alpha-band activity shows a similar pattern of fatigue/mind wandering levels across experimental factors: centro-parietal alpha-power increases over time-on-task and is elevated in the younger as compared to the older participants, while at the same time being sensitive to the motivational manipulation. Given this co-modulation with the fatigue/mind wandering scores, and the ample evidence that posterior alpha-power is inversely related to cortical excitability (Babu et al. 2018), we conclude that this increase in alpha-power is likely a marker of enhanced fatigue, rather than enhanced effort to maintain stable performance.

### Post-stimulus (task-related) oscillatory activity: two beta-signals that relate distinctively to response strategy and motivational manipulation

In analogy to the analysis of the pre-stimulus activity, we analysed post-stimulus oscillations in terms of effects of time-on-task first. We did this by identifying changes between experimental blocks 1 vs 8. To achieve this, we ran cluster-based statistics on the baseline-corrected time-frequency (TF) transformed data. We then examined their co-variation with age and the motivational manipulation. In a second step, due to the absence of motivational effects in the above analysis, we then tested directly for the effects of the motivational manipulation in the post-stimulus window (using cluster-based statistics comparing block 8 versus the motivational block 9 as a function of motivation group).

#### Post-stimulus beta-activity changes with time-on-task reflect qualitative differences in response strategy

Cluster-based permutation statistics between blocks 1 and 8 on post-stimulus TF-data identified a positive cluster in the lower beta frequency range (cluster statistic = 4899, p < .001) occurring in a late post-stimulus window (500-1000ms) (Figure 5A). The cluster is explained by a fronto-central beta-increase over time-on-task (detailed in Figure 5B).

**Figure 5.**
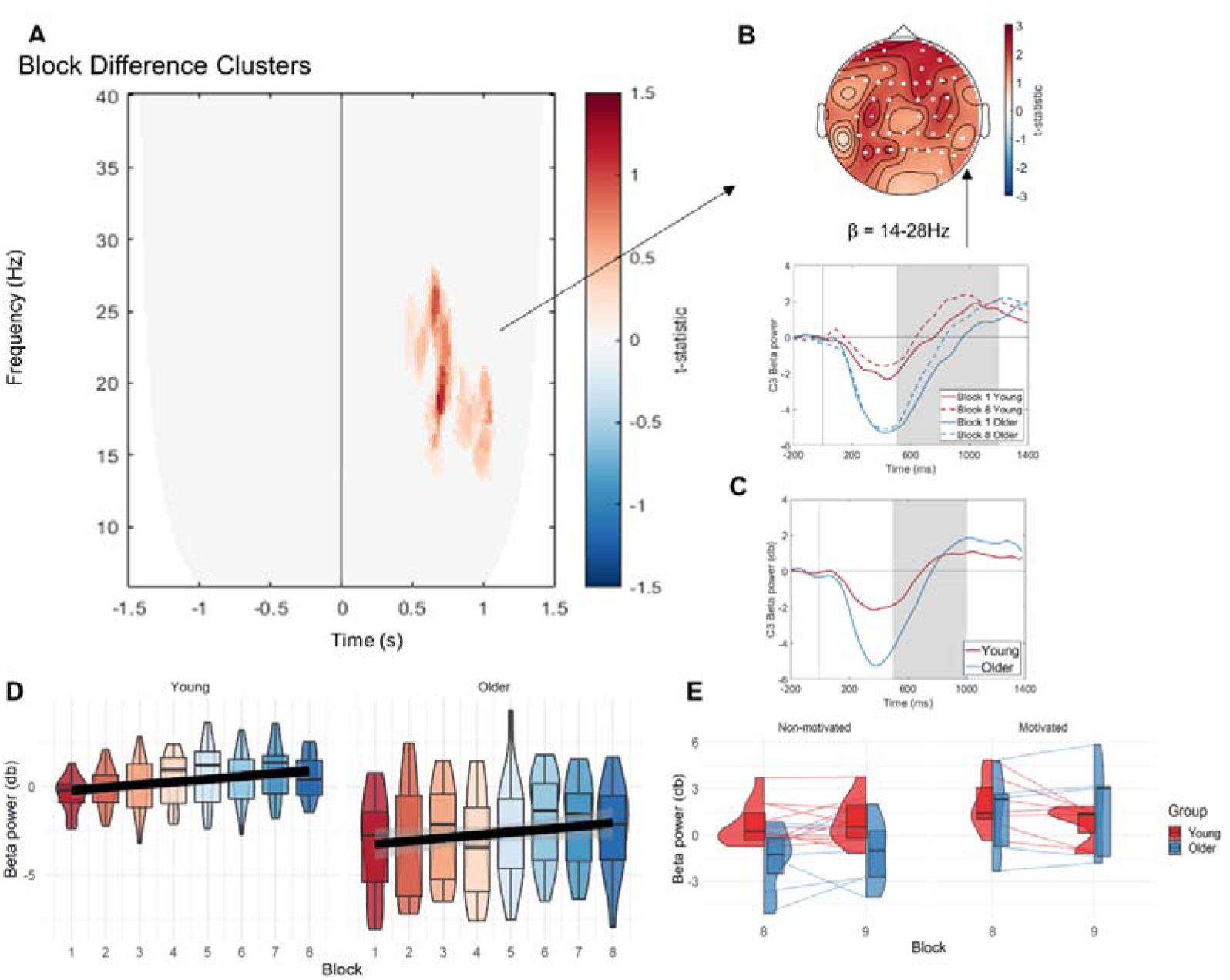
Time-on-task related oscillatory changes in the post-stimulus period. (A) Results of the cluster-based permutation test showing a time-on-task-related increase in the beta-band (14-24Hz) from block 1 to 8 in the later post-stimulus window (0.5-1s). (B) Topography of the time-on-task beta-change, indicating a fronto-central maxima. The line graph represents beta-changes across time-on-task and age-group illustrating that the positive cluster denotes an increase in beta-activity over time (compare solid vs dashed lines). (C) PSD comparison of young and older participants in the same frequency band across correct omission trial window with inferential time period area indicated. (D) Effect of time-on-task across all experimental blocks 1-8 per age-group (young, older), showing an overall positive trend independently of group, but higher beta-power in the younger participants. (E) No effects of motivational manipulation on the beta-signal, with no differential increase from block 8 to 9 between motivated and non-motivated participants, across the young (red) or older (blue) age group. Participant-level changes are depicted alongside median values and quartile ranges.

To examine beta-changes across all experimental blocks 1-8, we extracted this beta-signal (14-24Hz: relevant late time window) from the electrode with the most prominent cluster t-statistic (C3) per block and age-group (Figure 5D). A random mixed effects model with beta-power as the to-be-predicted variable confirmed its increase across time-on-task [t(32) = 2.98, β = 0.16, p = .005] and revealed a higher value in the young participant group [t(32) = -3.93, β = - 3.06, p < .001], but no interaction [t(32) = .147, β = 0.01, p = .884]. This indicates an overall linear increase of task-related beta-power across time-on-task that is independent of age-group, while showing overall group differences. It is tempting to interpret these differences in terms of an association with fatigue/mind wandering levels (in analogy to the pre-stimulus alpha-power changes). However, inspection of the event-related beta-change (Figure 5B, line plot) provides evidence for a relationship to behavioural response patterns and hence response strategy across age-groups instead.

Inspection of Figure 5B (line plots) reveals a prominent beta-desynchronization around response onset (mean reaction times were ∼400-500ms, see above), followed by a beta-rebound. The desynchronization was much stronger in amplitude in the older than younger participants, whereas the beta-rebound showed an earlier latency in the younger compared to the older group, reflecting the differences in their reaction times (see above). Given this dynamic pattern and the fronto-central topography, we interpret this beta-signal to reflect differences in motor response strategies between the groups. To further inform this interpretation, we explored to what extent this signal could be driven by the motor response. We therefore re-analyzed the beta-signal but taking into account correct omission trials only (hence eliminating any contamination by motor execution). Given the low number of omission trials, we averaged the data across all blocks 1-8 (it was not possible to resolve blocks 1 and 8 separately). This analysis revealed the same pattern (comparing Figures 5C vs 5B), including in terms of age-group differences (t(25) = -2.20, p = .037), which therefore suggests that this beta-signal is more of a cognitive or motor control nature than linked to motor execution. Based on these findings, we interpret the stronger beta-desynchronization in the older as compared to the younger group to reflect deployment of more effort towards accurate motor control, while we interpret the shorter latency in beta-rebound in the young, as compared to the older group, to reflect the speeded response strategy.

To test whether the time-on-task related beta-signal also co-varied with the motivational manipulation, this signal was extracted in the electrode with the highest t-statistic (C3) also for the motivational block 9 and separately for the motivated and non-motivated groups, for comparison with block 8 (see Figure 5E). Using a 2x2x2 ANOVA comparing motivation group (motivated, non-motivated), age-group (younger, older), and block (block 8 vs motivational block 9), we observed an effect of motivation group [F(1, 26) = 4.96, p = .035, η² = .142] showing that the group randomly allocated into the motivational condition had an overall higher beta-signal (Figure 5E). There was no main effect of block [F(1, 26) = 0.02, p = .889, η² < .001] or age group [F(1, 26) = 3.11, p = .089, η² = .094], nor were there any significant interactions. This therefore indicates that in contrast to the pre-stimulus alpha-signal, this post-stimulus beta-signal is not amenable to manipulation by motivation.

Overall, these analyses suggest that the qualitative differences in response strategy (accurate vs fast) across the age-groups are reflected in a beta-signal of fronto-central topography, which in its rebound-component is modulated with time-on-task as well as reaction times (see analyses below).

#### Motivation effects on beta-activity

Because we did not observe the time-on-task related beta-pattern to be altered by motivational manipulation (as per above), we tested for effects of motivation in post-stimulus (task-related) activity directly through cluster-based permutation tests. We compared TF-differences between blocks 8 and 9 across the two motivational groups (motivated vs. non-motivated) collapsing across both age-groups (interaction of motivation x block in TF-space). The analysis revealed a broad, late beta-cluster (Figure 6A), showing a weaker left fronto-parietal beta-increase from block 8 to 9 in the motivated relative to the non-motivated group (statistic = 351.87, p = .002; see Figure 6B for topography and time course).

**Figure 6.**
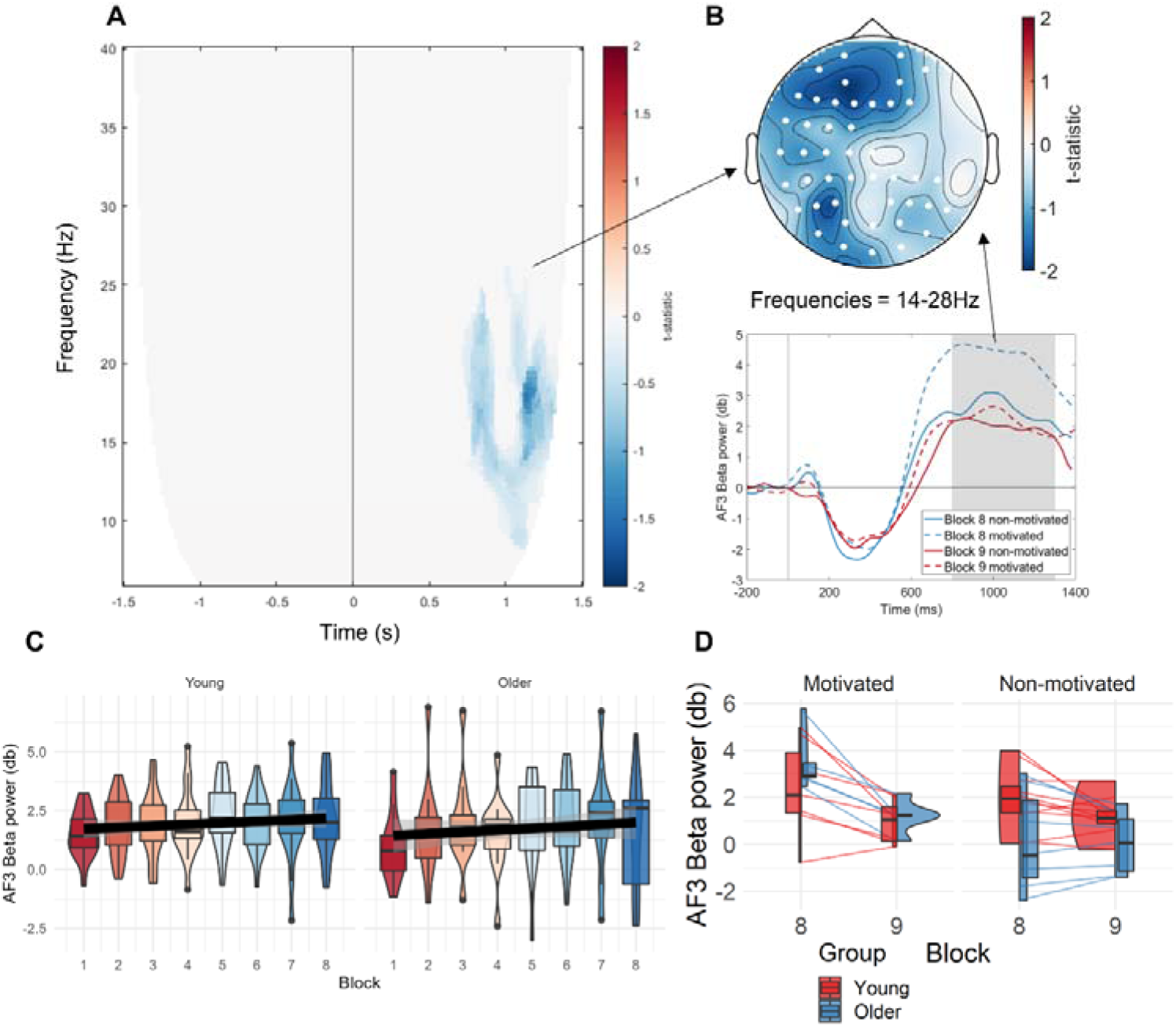
Motivation-related beta changes. (A) Results of the cluster-based permutation statistics testing the interaction motivation x block in TF-space (contrast of the difference signal of block (8 vs 9) between motivational groups). The results show a late (0.7 -1.2s) effect of motivational manipulation on beta activity (14 – 28Hz). (B)Topography and temporal dynamics of the motivation-related beta-change in the post-stimulus window indicating a left fronto-parietal decrease in beta-activity between block 8 to 9 in the highly motivated (dashed lines) but not the non-motivated participants (solid lines). (C) Evolution of the motivational beta-signal over time-on-task (blocks 1-8) per age group (young, older). The plot shows that this beta-signal is stable over time-on-task. (D) Effects of the motivational manipulation on the beta-signal across all groups and relevant blocks, illustrating a larger beta-decrease from block 8 to 9 in the motivated relative to the non-motivated participants independent of age-group.

To examine whether this signal also modulates with time-on-task and/or age-group, we extracted the motivational beta-signal (14-28Hz: in the relevant window) across all experimental blocks 1-8 and both age-groups from the electrode with the most prominent cluster t-statistic (AF3) in the test of motivation effects (Figure 6C). A linear model predicting this beta-power from across blocks and age groups showed no effect of either age [t(28) = -.652, β = -0.31, p = .520] or block [t(28) = 1.31, β = 0.06, p = .201], nor their interaction [t(28) = .234, β = 0.02, p = .817], indicating that this motivational beta-signal was unaffected by time-on task and age-group and hence distinct from the time-on-task beta-effect related to response strategy.

Finally, to test the beta-effects of motivation across all groups/conditions (Figure 6D), we run a 2x2x2 ANOVA with the factors motivation group (motivated, non-motivated), age-group (younger, older) and block (block 8, motivational block 9). The model revealed that the motivated group had higher beta-signals than the non-motivated group [F(1, 26) = 4.46, p = .045, η² = .121] and that there was a main effect of block[F(1, 26) = 21.36, p < .001, η² = .137] with a lower signal in block 9, but no effect of age [F(1, 26) = 1.71, p = .203, η² = .050]. The interaction between motivation and block was significant [F(2, 26) = 8.55, p < .007, η² = .060] independently of age-group (non-significant 3-way interaction motivation group x age group x block: F(3, 26) = 2.51, p = .125]). Post-hoc analysis exploring the interaction of motivation x block using dependent sample t-tests revealed that while the signal of the non-motivated group did not differ between the two blocks [t(17) = 1.94, p = .069], the motivated participants’ beta synchronisation decreased [t(11) = 4.45, p < .001]. There was also a significant interaction between age and motivation [F(2, 26) = 5.28, p = .030, η² = .141] but this interaction was not further explored, as it almost certainly reflects a chance finding (despite random allocation in age and motivation groups). No other interaction was significant.

Overall, the analyses suggest that motivation *per se* is reflected in a post-stimulus beta-signal of fronto-parietal distriubtion.

### ERP analyses

During the examination of the task-related signal, another cluster (statistic = 4577, P < .001) occurred in the early time window (0ms – 450ms), indicating a decrease in the alpha oscillatory band with the progress of task. As the cluster reflected early task-related alpha desynchronisation, it was further investigated for its possible connection to the P300 ERP component (Picton 1992; Studenova et al. 2023).

ERPs from the same time window of the signal coming from the highest un-lateralised electrode, detected as significant by the cluster permutation (Fz), were compared in their group peak maxima. A 2x2 ANOVA was conducted to examine the effects of block (1, 8) and group (younger, older), and their interaction on the baseline-corrected signal. The resulting model was not significant, F(3, 64) =.555, p = .647. Furthermore, the same signal was also traced into the motivational condition, compared in a 2x2 ANOVA collapsed across age, examining the effect of block (8, 9) and motivation (non-motivated, motivated). The resulting model was significant F(3, 56) = 3.47, p = .022. There was no effect of block (β = -0.21, t = -.396, p = .694), but the young group had a higher P300 amplitude (β = -1.24, t = -2.06, p = .044), with no interaction (β = -0.19, t = -.226, p = .822). A test was also run to detect an effect of trial type (commission, omission) and age group (young, older) on the P300 amplitude. The overall model was significant F(3, 64) = 7.70, p < .001. The test revealed no effect of trial type (β = -.162, t = -.216, p = .830), a higher P300 amplitude in young participants (β = 2.41, t = 3.31 p = .002) and no interaction (β = 0.130, t = .126, p = .900).

In summary, analyses of the post-stimulus oscillatory activity revealed two distinguishable beta-signatures: one that had a fronto-central topography and in its rebound-component was modulated with time-on-task, as well as reaction times, as a likely marker of response strategy, and another of fronto-parietal distribution modulated by motivation *per se*. The P300 did not reflect any of our manipulations, just higher amplitudes in the younger compared to the older age group.

### Relationship between EEG signals and behaviour

To detect links between behaviour, namely measures of subjective state fatigue, mind wandering and reaction time, and the neural markers of time-on-task, age effects and motivational manipulations (see frequency bands/electrodes identified in the above EEG analyses), we built multiple linear regression models predicting subjective and RT measures by neural signals with addition of the effect of age and their interaction. Single electrodes with the highest t-statistic resulting from permutation tests were extracted and changes in signal were compared to changes in subjective measures.

The model testing state fatigue (VAS) changes was not significant for its relationship to pre-stimulus alpha-changes (electrode CP1) [F(3,30) = 2.25, p = .103], or task-related beta-changes (electrode C3) as extracted in blocks 1 and 8 [F(3,30) = .774, p = .518]. Likewise, no effect was found for the relationship between mind wandering to pre-stimulus periodic alpha change [F(3, 30) = 1.73, p = .182], or task-related beta [F(3, 30) = 2.43, p = .085].

The model testing reaction time change was not significant for its relationship to pre-stimulus alpha-change (electrode CP1) [F (3,30) = 1.54, p = .224], but the model was significant for task-related beta-change (electrode C3) [F(3,30) = 8.84, p < 0.001]. There was no main effect of beta-signal [β =.004, t = .224, p = .824], but a significant interaction between beta signal change and age [β = - 0.06, t = -3.09, p = .004]. An inspection of Fig 8 reveals that the relationship between beta change and reaction time change was driven by the older adult group. Older adults who showed a greater decrease in reaction times, showed a greater increase in beta power. No effect was found for the relationship of error change to pre-stimulus periodic signal change [F(3, 30) = 0.192, p = .897], or task-related beta [F(3, 30) =.185, p = .906]. The relationship between behavioural change and motivational beta-signal difference between blocks 1 and 8 was not tested as the previous model showed no modulation of the signal with either age or time-on-task.

**Figure 8.**
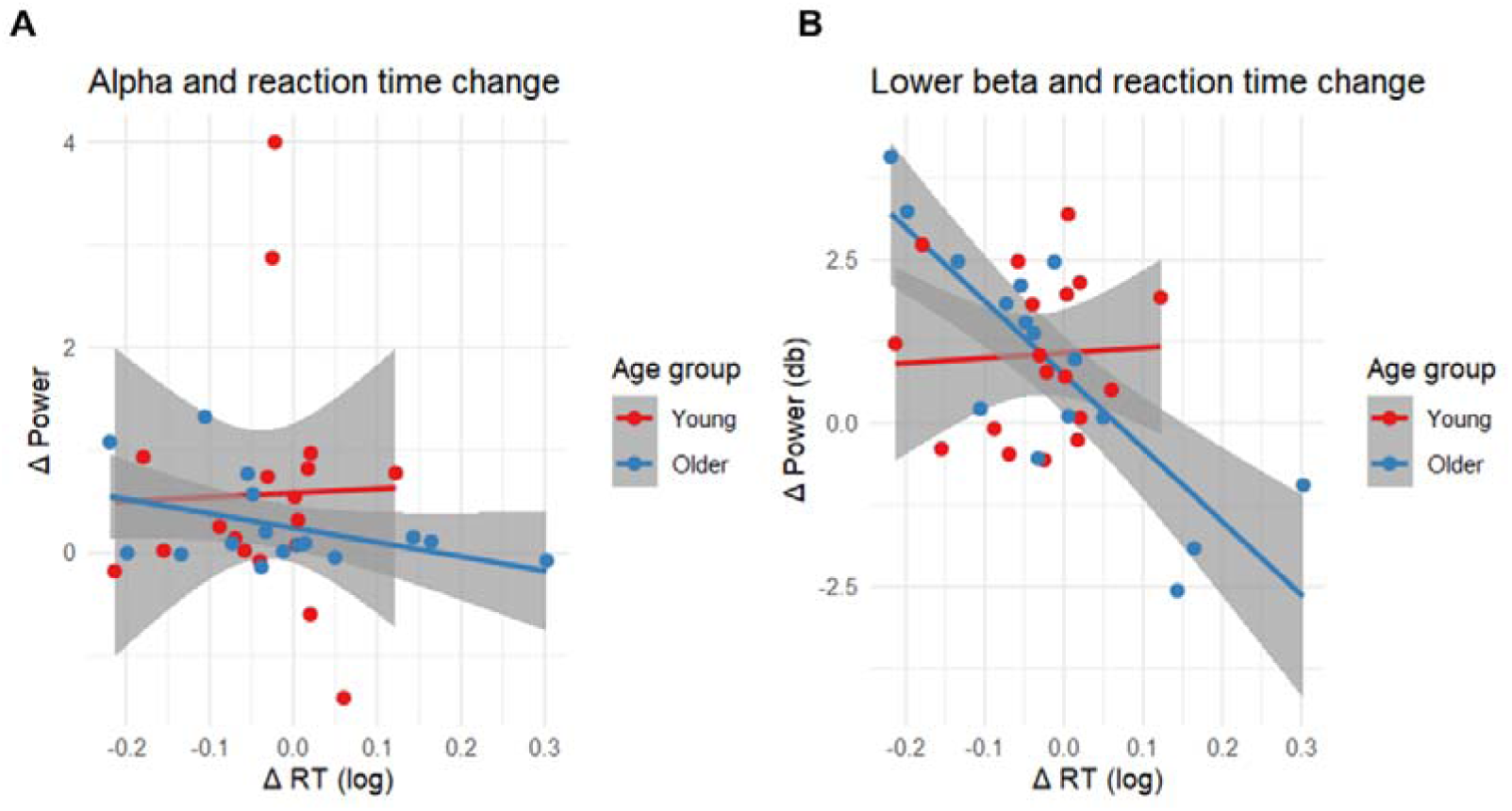
Brain and behaviour link. Relationship between the change in reaction times and (A) the change in alpha power over CP1, (B) change in lower beta power over C3 across the main experiment over time-on-task for both age groups.

## Discussion

This study sought to investigate and identify neural patterns underlying vigilance, fatigue and motivation during sustained attention. Its results provide new evidence for three distinct oscillatory patterns associated with fatigue and motivation and unexpectedly response strategy.

### Pre-stimulus Alpha Oscillations

The task elicited a distinct linear increase in pre-stimulus alpha synchronisation over time. This time-on-task rise reflects other research around time-related changes in EEG signal arising from experimental manipulation (Tian et al. 2018; Li et al. 2020; Jacquet et al. 2021) and adds to the existing body of literature highlighting detectable changes in alpha oscillations in relation to demanding tasks (Benwell et al. 2019; Huycke et al. 2021; Pershin et al. 2023). More importantly though, our findings link these centro-parietal alpha-power increases to a general rise in subjective fatigue and mind-wandering, as well as the motivational manipulation: not only was there as a significant alpha rise over time (blocks 1 to 8), but also significantly greater elevation of the signal in the younger compared to the older participants. The signal thus mirrored perfectly, the reported fatigue/mind wandering scores which were not only significantly elevated over time, but also significantly higher in the younger compared to the older participants. Finally, the signal was co-modulated with participant motivation: while alpha oscillations kept rising for the unmotivated participants of the final block (in line with the reaction times that also increased from block 8 to 9 in this group), it levelled out for the motivated group (who were promised a monetary incentive if they outperformed others).

As a new finding and interpretation, in particular in light of the reported absence of a behavioural vigilance decrement, we propose this centro-parietal pre-stimulus alpha increase during time-on-task, that further levels out with participant motivation, to not reflect a resource depletion (nor a disengagement of attention (Arnau et al. 2021; Krigolson et al. 2021), but instead to closely mirror the waxing and waning of a subjective fatigue state.

### Post-stimulus, time-on-task modulated beta-oscillation relate to response strategy

We also observed a time-on-task rise in fronto-central post-stimulus beta synchronisation which, in contrast to the alpha oscillation, was unaffected by motivational interference. The observed pattern followed the structure of a classic post-motor beta desynchronization and rebound, the specific neural response concerned with preparation (Swann et al. 2009; Darch et al. 2020) and/or execution of a motor response (Parkes et al. 2006; Heinrichs-Graham et al. 2017). In the present study, the desynchronisation was much stronger in the older compared to the younger group, whereas the beta-rebound showed an earlier latency in the younger group. Given this dynamic pattern and the fronto-central topography, we interpret this beta-signal to reflect age differences in motor response strategy. More specifically, we think that these age (strategy) differences are of a cognitive or motor control (rather than motor execution) nature as we found the same pattern also for omission trials (which lacked a motor execution aspect). Our thinking is further underpinned by the significant relationship we found between beta signal change and age: older adults who showed a greater decrease in reaction time over time-on task, also showed a greater increase in beta power. This pattern was absent in the younger group. Overall, these analyses suggest that, across age groups, there are qualitive differences in response strategy (accurate vs fast), that are reflected in a beta-signal of fronto-central topography: we interpret the stronger beta-desynchronization in the older, as compared to the younger group, to reflect a deployment of greater effort towards accurate motor control, with the shorter latency in the beta-rebound in the younger group, to reflect the speeded response strategy. These results further emphasise contrasts in age group specific response strategies reported previously (Lara et al. 2014; Dang et al. 2018; Reteig et al. 2019; Statsenko et al. 2020; Vallesi et al. 2021), but more importantly, we now demonstrate that these qualitative strategic differences are underpinned by distinct beta oscillations (see (Xifra-Porxas et al. 2019) for similar findings on grip strength).

### Post-stimulus ‘motivational’ beta oscillation

In addition to the beta-signature with its fronto-central topography, we found another beta oscillation of fronto-parietal distribution that was independent of age-group but instead modulated by motivation *per se*. This signal was stable over time-on task, yet we found it to show a larger beta-decrease from block 8 to 9 in the motivated relative to the non-motivated participants. This signal decrease was coupled with a reaction time levelling out in the motivated group (from block 8 to 9), while reaction times in the non-motivated group increased significantly. Our findings thus show that the motivation manipulation was effective, similar to the results of Reteig et al. (2019) who showed a temporary increase in sustained attention task performance after an (unexpected) motivational manipulation. Most importantly, we show this motivational effect to be co-modulated with an oscillatory beta decrease, whereas Reteig and colleagues failed to find any EEG markers linked to their motivation manipulation. Beta-oscillatory changes have previously been linked to motivational initiative (Wilhelm et al. 2022) and reacted to changes of internal state also (Nickel et al. 2020), although there have been no previous reports on attentional motivation manipulation in human participants.

Our data thus best reflect the findings and interpretations of Stoll et al., (2016) who examined modulations of frontal beta around spontaneous pauses in work in monkeys. They found that after pauses, the beta power modulation would reset, and the cognitive control effect (task performance) maintained. We report this signal resetting and maintenance of performance not for pauses in work, but instead for our motivation manipulation. In fact in line with our data, Stoll and colleagues (2016) propose that frontal beta oscillations reflect multiple factors contributing to the regulation of cognitive control and that motivation parameters can act as modulators of cognitive control.

## Limitations and future work

Our study reproduced many typical age related findings, including higher baseline mind wandering and lower reaction times (Learmonth et al. 2017; Fountain-Zaragoza et al. 2018; Diede et al. 2022), as well as strategy differences ((Vallesi et al. 2021) prioritising speed over accuracy (Lara et al. 2014) in the younger over the older age group. We also found typical age characteristics in the pre-stimulus oscillatory window regarding the structure of the aperiodic components of the signal in particular, with steeper exponents and higher intercepts (Donoghue et al. 2022; Turner et al. 2023) for the younger compared to the older age groups.

What was surprising was the absence of a decisive vigilance decrement, contrary to our hypothesis and in contrast to other work (Kaufman et al. 2016; Gartenberg et al. 2018; Walker and Trick 2018; Reteig et al. 2019; Pershin et al. 2023). The speeding up of reaction times when taking participant-level randomised intercepts into account, replicates our own earlier behavioural findings (Hanzal et al. 2024) with a small effect of task time on speed (blocks 1 to 8). As the participant accuracy did not show a corresponding decline, this result cannot be interpreted as a vigilance decrement. Having consulted previous work (Staub et al. 2014; Reteig et al. 2019) where a decrement occurred at 20-35 minutes that continued being present in the following hour, we too expected a decline in our study, where the actual task took 45 minutes, and think that a longer experimental duration would have led to an eventual lapse of this maintained performance (Martínez-Pérez et al. 2023).

Unfortunately also, despite the clear co-modulation of the reported alpha rise over time with fatigue and mind-wandering, and with these rises also closely mirroring the age (elevated tiredness in the younger) and motivation (reaction time rises in the unmotivated group (blocks 8 to 9)) manipulation, we did not find a statistically significant correlation between the reported alpha rises and the reported increases in fatigue and mind-wandering. It is possible that the subjective fatigue and mind wandering scales we used lacked sensitivity and further work is needed here. We would argue that our findings are a first pointer to neural changes as better and more sensitive markers of fatigue than subjective reports, but new studies are needed to further firm up that centro-parietal alpha-power increases reflect increases in fatigue.

Finally regarding the P300, the only effects we found were higher amplitudes in the younger over the older age group, yet we failed to detect the established task-related pattern of an occipito-parietal P300 amplitude decrease, associated with either effortful processing (Hart et al. 2012) (Hart et al., 2012) or fatigue (Egner and Gruzelier 2004; Hart et al. 2012; Krigolson et al. 2021; Kustubayeva et al. 2022).

## Conclusion

We report that undergoing 45 minutes of a sustained attention task (SART) induced subjectively elevated fatigue and mind wandering scores that where co-modulated by a centro-parietal alpha power rise, suggesting pre-stimulus EEG oscillations as a possible sensitive, objective marker of fatigue. Post-stimulus activity revealed two distinguishable beta signatures: a fronto-central topography as a marker of time-on-task strategy and a fronto-parietal distribution modulated by motivation *per se*. We suggest that these two latter signals reflect a (motivational) cognitive control mechanism behind resetting a performance decrement that is independent of persistent fatigue.

